# Population coding strategies in human tactile afferents

**DOI:** 10.1101/2022.05.04.490609

**Authors:** Giulia Corniani, Miguel A Casal, Stefano Panzeri, Hannes P Saal

**Author notes:** Correspondence: SP; HPS. Co-first authors (Giulia Corniani, Miguel A Casal).

## Abstract

Sensory information is conveyed by populations of neurons, and coding strategies cannot always be deduced when considering individual neurons. Moreover, information coding depends on the number of neurons available and on the composition of the population when multiple classes with different response properties are available. Here, we study population coding in human tactile afferents by employing a recently developed simulator of mechanoreceptor firing activity. First, we demonstrate that the optimal afferent density for conveying maximal information depends on the tactile feature under consideration and the afferent class coding this feature. Second, we find that information is spread across different classes for all tactile features, such that combining information from multiple afferent classes improves information transmission, and is often more efficient than increasing the density of afferents from the same class. Finally, we test the importance of timing precision and afferent identity in the population code to probe whether temporal and spatial information can be traded against each other. Destroying temporal information turns out to be more destructive than removing spatial information, and the contribution of either cannot be completely recovered from the other. Overall, our results suggest that both optimal afferent innervation densities and the composition of the population depend in complex ways on the tactile features in question, potentially accounting for the variety in which tactile peripheral populations are assembled in different regions across the body.

## Introduction

The brain processes information and makes decisions based on the activity of a large number of neurons (1). Studying population activity can reveal aspects of the neural code that are obscured when only individual neurons are considered (2). For example, the well-known population vector technique has shown that the direction of arm movements can be precisely decoded from a population of cortical motor neurons, even though individual neurons are only broadly tuned to direction (3). Moreover, some coding strategies will become evident only if the responses of multiple neurons are considered. For example, while a neuron that remains silent to a certain stimulus might not appear to convey any information at all, when it is part of a larger population where other neurons are responding, this silence can be meaningful (4). Response correlations between neurons also affect decoding (see 5, for an example). Furthermore, populations often consist of heterogeneous classes of neurons, especially in sensory systems, such as the diversity of retinal ganglion cells in the visual pathway (6) or the different classes of tactile neurons in the somatosensory periphery (7). Theoretical studies have shown how response properties and class membership of individual neurons can be optimized to maximize joint information coding in the population (8–10). However, because this optimization relies on the full population, predicting how or to what extent an individual neuron contributes to population coding becomes impossible without considering the properties of other neurons that make up the population. Given these findings, it is thus paramount to study the population activity of sensory neurons in order to understand what stimulus information is available at subsequent processing stages.

Tactile interactions are mediated by mechanoreceptive afferents and the glabrous skin of the human hand is innervated by approximately 17,000 fibers (11). These are divided into different classes based on their response properties and receptive fields. Three classes are mainly involved in discriminative touch (SA1, RA, PC): SA1 afferents exhibit small receptive fields and respond to static or low-frequency indentations, RA afferents possess slightly larger receptive fields and respond to dynamic flutter stimuli, and PC afferents exhibit extremely large receptive fields and are most responsive to very high frequency vibrations. These classes also differ in the density with which they innervate the skin, both compared to each other and at different locations on the skin (11). A stimulus applied to a specific skin area will typically activate hundreds if not thousands of afferents of different classes all responding with distinct spiking responses (12). However, peripheral neurophysiological measurements are subject to technical limitations, and typically only one or a small number of afferents are recorded at once. Moreover, many studies place the stimulus directly above the targeted afferent’s receptive field hotspot, in an effort to maximize neural responses within the limited recording window, but such a setup implies that responses from receptors located away from the contact location will be neglected. Given these constraints, afferent activity on a population level has scarcely been investigated, and, consequently, our understanding of how tactile information is represented in the peripheral population is limited (though see 13, for a summary of tactile population codes).

A particular source of debate in the tactile literature has been the role of different afferent classes. Traditionally, each afferent class was thought to carry information about different and complementary stimulus features (14). However, more recently it has become clear that most natural stimuli elicit responses from multiple afferent classes simultaneously (see summary in 15), for example in texture perception (16). Furthermore, both experimental evidence (17) and computational modeling (18) suggest that information from multiple classes of afferents is integrated in cortex, if not before, and psychophysical studies have revealed that the quality of a tactile percept does not necessarily depend on receptor class (19). However, to what extent peripheral tactile population activity carries complementary information about relevant stimulus features in different afferent classes has not been quantified and it is therefore unclear when and how it would be beneficial to integrate such information.

Here, we investigate the contribution of large neural populations in tactile stimulus coding and examine the interplay of tactile submodalities in this process. Because the lack of population level data currently precludes empirical study, we used a large-scale computational model, Touchsim (20), to simulate the activity of hundreds of peripheral tactile afferents of three classes in response to naturalistic stimuli, similar to those commonly used in experimental settings. First, we parametrically studied the role of afferent density in single-class afferent populations to explore if and how the composition, and particularly the number of afferents, affects the stimulus information encoding. Secondly, we considered the three classes together and asked whether each class encoded complementary or redundant information regarding stimulus features. Finally, we assessed the importance of temporal and spatial encoding precision when considering afferents on a population level. Overall, our work demonstrates that a population-level view of tactile coding is crucial for a thorough understanding of tactile information processing.

## Results

We used a large-scale neural simulator (20) to simulate the spiking activity of individual afferents belonging to three afferent classes (SA1, RA, PC) jointly spanning the range of tactile sensitivity. In our setup, we simulated the responses of a population of receptors placed along a line extending outwards from the contact location of the stimulus probe (see Figure 1A). This spatial arrangement of receptors allowed for systematic manipulation of the receptor density in the simulations. Sixteen different afferent populations with a density ranging between 1 and 140 afferents/cm^2^ were considered for each afferent class. Therefore, we could test the effect of low, medium, and high densities (Figure 1B) on information encoding and also directly examine natural innervation densities, such as those encountered on the palm or finger (Figure 1C). The simulated stimulus was a circular probe indented into the skin and then vibrated. We varied four stimulus features systematically across trials: the probe size (1-4 mm), the depth of the indentation (0.3-1.2 mm), the ramp length (10-50 ms), and the vibration frequency (0-200 Hz) (see Figure 1D and Methods). These parameters were chosen to span the range of tactile stimuli that are typically experienced. They are also similar to stimuli commonly employed in neurophysiological experiments, such as those used to fit the initial Touchsim model (20), and simulated responses can therefore be expected to be a close match to what would be recorded in an actual experiment. Finally, varying the stimulus across multiple parameters simultaneously ensures that the complexity of everyday tactile interactions is reproduced in the resulting population responses.

**Fig. 1.**
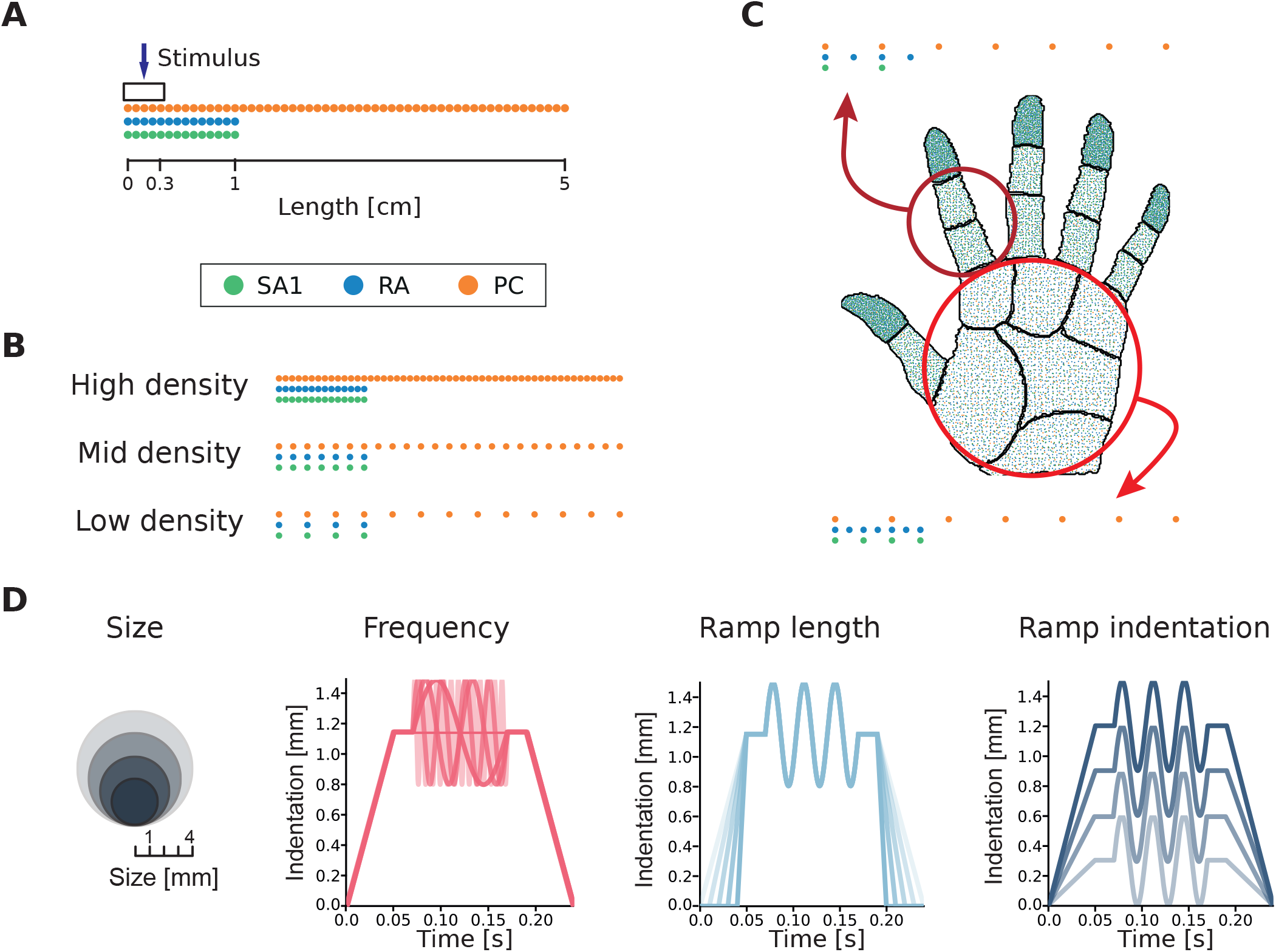
Simulation setup. **(A)** Example of afferents terminating along a line, radiating outwards from the probe centre (indicated by the arrow). The probe has circular shape (of varying size) and is centred on the origin of the line. Dots of different colors corresponds to different afferent classes (separated in the illustration to facilitate visualization). **(B)** Example of afferent populations with different densities. **(C)** Representation of afferent densities measured on the human hand and corresponding simulated populations distributed over a line that mimic the densities observed in the palm and finger. **(D)** Illustration of the different stimulus features considered: probe size, vibration frequency, ramp length, and ramp indentation depth.

To analyse the simulated responses, we coupled advanced machine learning techniques with information-theoretic analysis to compute how much information about each stimulus feature was encoded in the activity of different populations of afferents (see Methods for details). In short, after simulating the spiking responses (Figure 2A), we first used Non-Negative Matrix Factorization (NMF; 21) to succinctly capture the spatiotemporal patterns of neural responses for each afferent class (Figure 2B). This technique linearly decomposes each single-trial spatiotemporal sequence of spike trains into a sum of non-negative spatiotemporal modules (describing the recurrent spatiotemporal patterns of firing of the population) and non-negative activation coefficients (describing how strongly each pattern is recruited in a given trial). NMF was chosen to reduce the dimensionality of population activity because it is a natural decomposition for spike trains, which are by definition non-negative, because it can give accurate single-trial representations of activity even when neural responses are non-orthogonal and overlapping from trial to trial, and because its coefficients are biologically interpretable (22, 23). Following previous work (24), we then approximated the probabilities of occurrence of the NMF activation coefficients using a Generalized Linear Model (GLM; see 2C). We then used this probabilistic model to compute the posterior probability of each stimulus feature given the observation of the spatiotemporal population spike train in each trial. Finally, we computed the information using the posterior probabilities between the presented and the decoded stimulus (2D). This procedure provides a data-robust but effective lower bound to the total information carried by population activity (24).

**Fig. 2.**
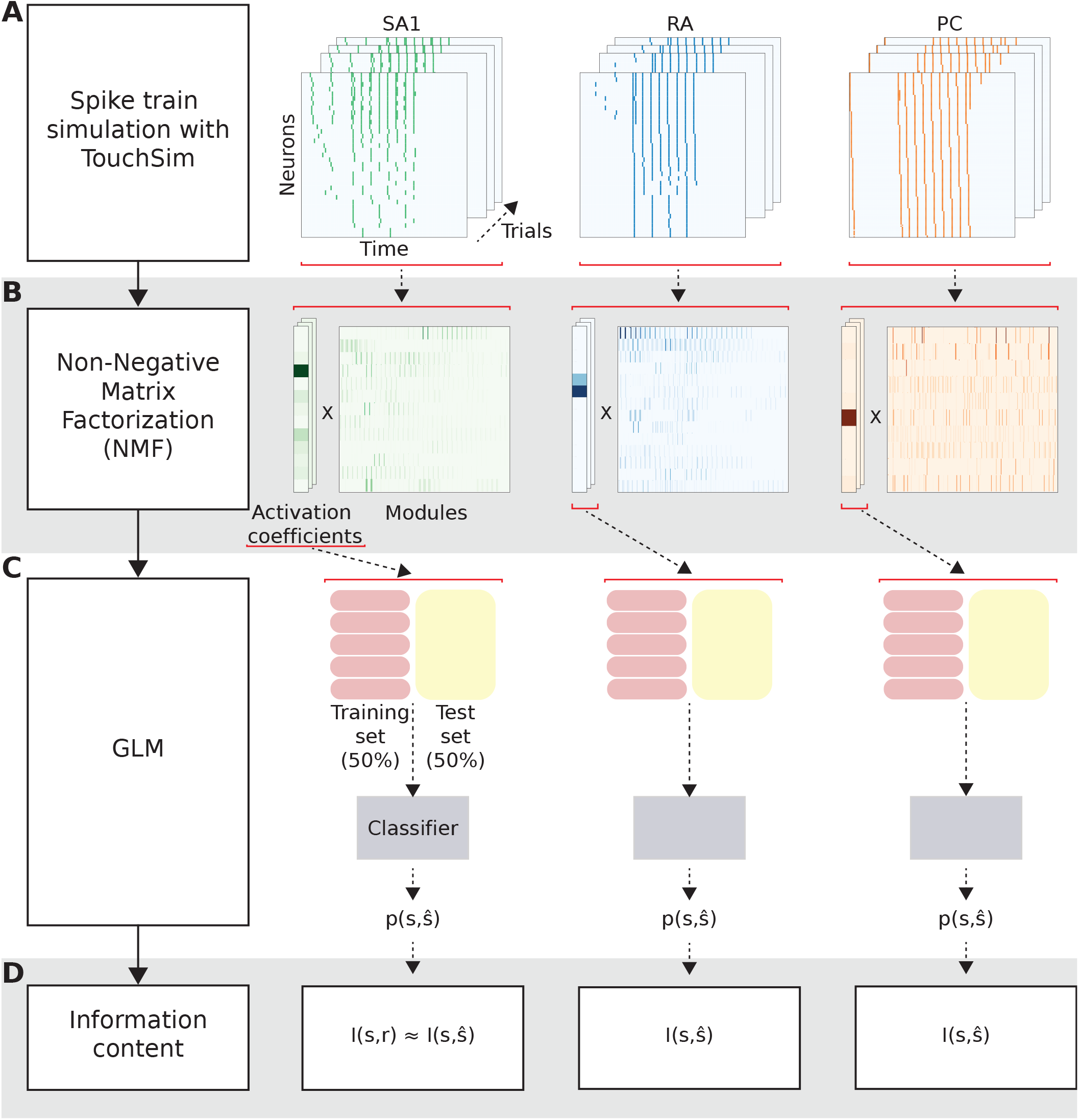
Analysis pipeline and calculation of information. **(A)** The spike trains are simulated using Touchsim. **(B)** The spike matrices are then decomposed using the Non-Negative Matrix Factorization (NMF) method, obtaining a set of non-negative activation coefficients and modules. **(C)** A Generalized Linear Model (GLM) fed with the neural activity captured in the NMF activation coefficients gives the probability of observing each stimulus feature. **(D)** Probabilities are used to compute mutual information (MI), representing the information that the neural activity carries about the stimulus.

### Information carried by individual afferent populations

In a first analysis, we investigated the information carried by each of the three afferent populations separately. To understand which afferent population best encoded any given feature and how the information depended on the spatial density of the afferents, we calculated the total information carried by each population (Figure 3A) by simulating responses with different spatial receptor densities.

**Fig. 3.**
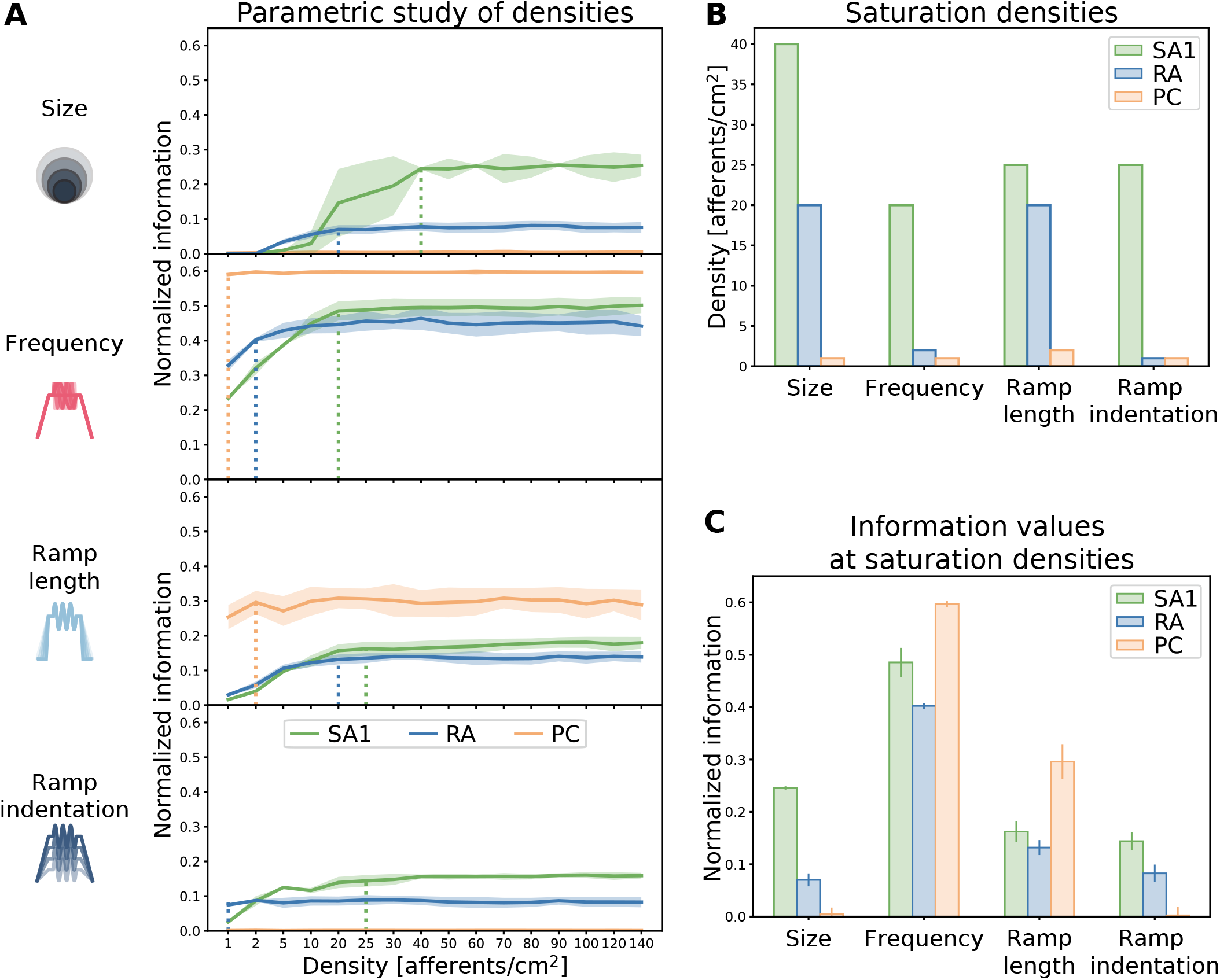
Effect of afferent density on stimulus feature coding. **(A)** Information content (normalized by the stimulus entropy) for different stimulus features provided by single-class afferent populations of varying density. Solid lines represent the average over 40 trials, shaded regions represent standard deviation. Dotted vertical lines indicate information saturation points. **(B)** Saturation densities for each feature and afferent class. **(C)** Maximum information content provided by each afferent class at the saturation density for each feature.

For encoding stimulus size, we found that SA1 afferents were most informative, with information increasing and then saturating at a density of 40 afferents/cm^2^. RA afferents provided more information at very low densities and saturated at a lower level (20 afferents/cm^2^). In contrast, PC afferents did not carry any information about stimulus size at any of the densities considered. This result can be explained by the fact that PC afferents exhibit extremely large receptive fields (25), certainly larger than the differences in size between the stimuli we applied.

Next, we considered the encoding of the frequency of stimulation. PC afferents provided the highest frequency information, as predicted by the fact that PC afferents are well known to carry frequency information in vibrotactile stimulation (14). Given their large receptive field size, frequency information of PC cells already saturated with the lowest density of afferents considered. In agreement with previous studies, RA afferents also carried considerable information about frequency (14). SA1 populations carried low amounts of frequency information at small spatial densities, but slightly exceeded the frequency information of RA afferents at higher spatial densities. This result may appear to contradict earlier empirical studies, where SA1 afferents were shown to respond only to the lower extreme of the range examined in our study (26). However, in our simulations the sinusoidal wave is superimposed on a ramp-and-hold indentation. This sustained indentation causes low spiking activity in the SA1 afferents, with spikes aligned to the vibration (see Figure S1B). Our finding suggests that this information emerges when taking into account the activity of SA1 afferents on a population level rather than single afferents separately.

PC afferents were also the most informative class about the stimulus ramp length, followed by SA1 and RA afferents, which provided similar levels of information, but required higher densities than PCs to reach saturation. Finally, SA1 afferents carried the highest amounts of information about ramp indentation, with PC afferents not encoding any information. RA afferents again provided higher information than SA1 ones at the lowest density, but adding more fibers did not increase information for this class.

The information saturation density, which we defined as the smallest value of density at which the population carried the asymptotic value of information reached for the highest simulated density, was the highest across classes for SA1 afferent for all considered stimulus features. Conversely, the information saturation density was the smallest for PC afferents in all cases (Figure 3B). Notably, when considering purely spatial features such as the stimulus size, the population encoding the highest asymptotic information level corresponds to the one with the highest saturation density (Figure 3B and C). Consequently, a high density of afferents is required to extensively innervate a skin area and discriminate between fine differences in the shape of stimulation. On the other hand, when looking at temporal features such as the frequency or the ramp length, sparsely distributed PC afferents overcome the information content encoded by the other more densely packed afferents classes.

Finally, our result shows that the RA class at saturation density always encodes less information than the SA1 and PC populations about any feature considered in this study (Figure 3C). However, at low densities (<10 afferents/cm^2^), RA afferents were more informative than SA1 for all features considered, suggesting that the optimal way to encode a tactile feature might depend on the number of neurons available.

### Information encoded by multiple afferent classes

Next, we investigated how tactile stimulus information was encoded in the joint activity of multiple afferent classes. In particular, we asked whether the information about stimulus features carried by an afferent class adds to and complements the information carried by other classes or whether the information carried by different afferent classes is redundant. To answer this question, we computed the information carried about each stimulus feature by the joint activity of populations of two or three afferent classes and compared the resulting values with the single-class information calculated above. Specifically, we used the concept of complementary information (27): we defined the complementary information carried by additional afferent classes over that of a reference class as the information carried by all considered classes jointly subtracted by the information carried by the reference class alone. All possible combinations of classes were considered. In these calculations, unless otherwise stated, we set the density of each class to the one measured on the glabrous skin of the human finger (see Methods for details). This allowed us to compare the information contribution of different classes in a realistic and biologically relevant setting.

We first considered whether afferent classes that were not the principal source of information about a stimulus feature added information that was complementary to that of the principally contributing afferent class. To do so, for each feature, we quantified the amount of complementary information that the less informative classes add to the information carried by the most informative class (Figure 4A). The amount of this complementary information was normalized to the amount of stimulus information carried by the most informative class. For all features, we found that the second and third most informative classes added information that complements the information carried by the most informative class alone. On average, the second most informative class added between 12 and 25% complementary information, depending on the feature considered. When considered jointly, the second and third most informative classes added on average between 15 and 30% of complementary information, compared to the most informative class alone. This result indicates that for each tactile stimulus feature, each class encodes some amount of complementary information about the stimulus that cannot be found in the activity of the other two classes.

**Fig. 4.**
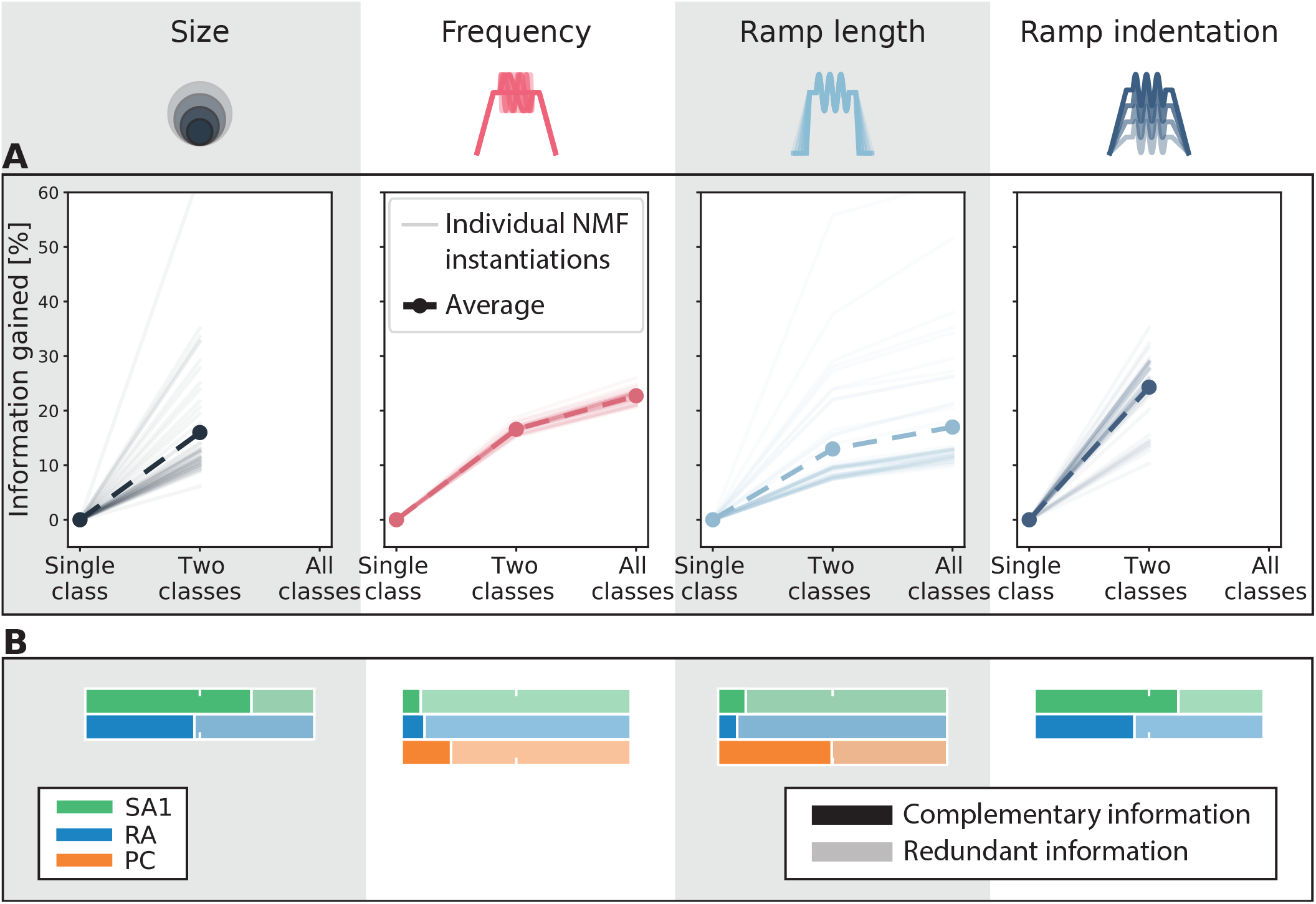
**(A)** Information gain considering the two and the three most informative classes together with respect to the most informative class alone. The density measured on the human finger was taken for each afferent class. Thin lines correspond to different instantiations of the NMF decomposition, and the thick dashed lines correspond to their averages. **(B)** Decomposition of information into redundant and complementary contributions of each class with respect to the remaining two classes together. Information for each afferent class has been normalized to 100% and was calculated at the density measured on the human finger. Note that in both panels **(A)** and **(B)** only two afferent classes were considered for the analyses regarding stimulus size and ramp indentation since the third class (PC) was carrying null information (see Figure 3C).

Next, we investigated which amount of each afferent class’s information contribution was complementary or redundant when considered against the information contribution of the other afferent classes. For each stimulus feature and each individual afferent type, we computed the fraction of the information carried by the considered afferent that is complementary with respect to the information already carried by the other two afferent classes. This fraction is an index of the specific novelty of the information of a given afferent class with respect to all others (Figure 4B). In general, a significant fraction of information carried by each afferent class was complementary to that of other classes. In most cases however, this fraction was not close to 1, meaning that there was also redundancy between the information carried by afferent classes. When examining how this fraction varied across stimulus features, interesting patterns emerged.

For vibration frequency and ramp length, the two stimulus features for which all three afferent classes encoded considerable information, we found mostly redundant coding, with relatively small fractions of complementary information (on average 13% for frequency and 23% for ramp length, Figure 4B). All three classes encode vibratory stimuli by locking their spiking activity to the sinusoidal traces, which explains the redundancy across classes. However, the fact that the frequency ranges encoded by each class do not completely overlap explains the existence of significant fractions of complementary information across all three afferent classes. Given that all three classes encoded large amounts of frequency information, the actual amount of complementary information added by each class was surprisingly large (Figure 4A). A similar pattern of complementarity and redundancy of information was observed for ramp length, which like frequency is a dynamic feature that depends on timing.

For stimulus size, the SA1 afferent population carried most of the information (Figure 3), and this information had a high value of complementarity (72%), indicating that it could not be found in other afferent types (Figure 4B). The RA afferent population added less information (Figure 3), but also exhibited a relatively large fraction of complementarity (48%) (Figure 4B). The encoding of size for SA1 and RA afferents seems to depend on the number of afferents that are activated by the stimulus (Figure S1A), and the observed complementarity between RA and SA1 afferents is partly due to differences in spatial sensitivity across the two populations. Information carried by PCs about probe size was negligible and therefore this class was not considered in the complementarity analysis for this feature.

Finally, for ramp indentation we found results that resemble those for stimulus size. The SA1 population carried most information, which was largely complementary (63%) to that of other classes. RA afferents carried less information than SA1 afferents, but part of this information (43%) was complementary to that of SA1 afferents. In this case, the encoding appears again to depend on the fraction of afferents that are activated by the stimulus, as was the case for stimulus size. This is a genuine form of population coding that would not be evident from single afferent analyses. PCs again provided negligible information (see Figure 3C), and thus were not taken into account.

### Effect of afferent density on complementary information

Having established that information about individual stimulus features is carried by multiple, rather than single, afferent classes and that different afferent classes often carry complementary stimulus information, we next asked how the complementarity of information depends on the spatial density of afferents. We were especially interested in whether, given the functional properties of afferents in each class, it would be more efficient to allocate all receptors to the most informative class or spread the receptors across different classes to take advantage of the complementarity of different classes. To address these questions, we systematically analyzed the information carried by individual afferent classes and their combination at different densities (Figure 5A). We tested the same upper and lower density limits as used previously. For a more realistic comparison with human biology, we also considered two other cases of spatial density arrangements, in which each population has a density equal to that experimentally found either in the palm or in the finger of the human hand (see Methods for precise numbers).

**Fig. 5.**
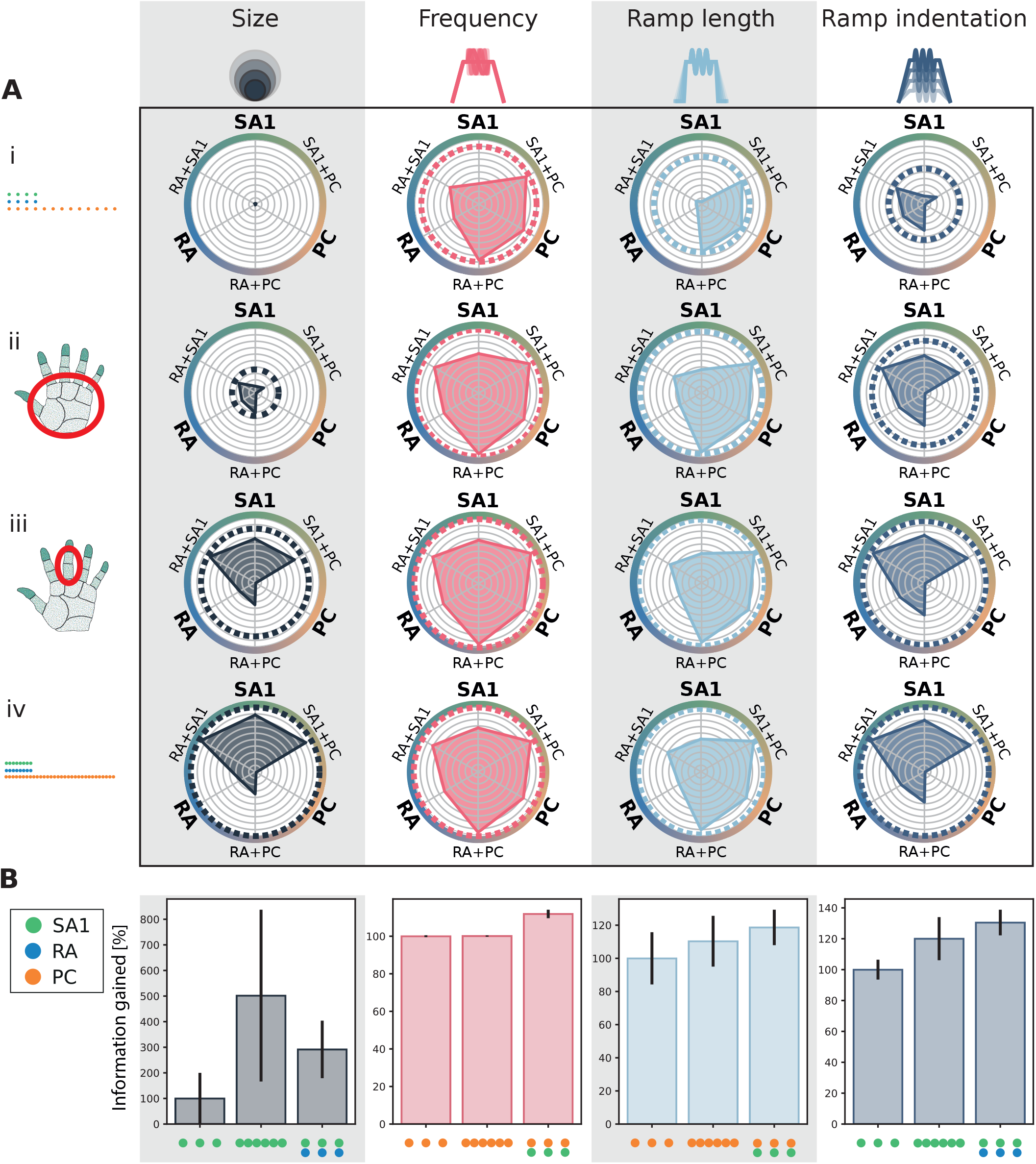
**(A)** Radar plot of the information content provided by single and combined classes at different densities for individual tactile features. Each radial axis represents the information content of a single afferent class or combination of two classes. Dotted circular lines correspond to the information given by the three afferent classes together. Information is normalized for each stimulus feature with respect to the information provided by the three classes altogether at the upper-limit density (i). Four different density sets were considered: (i) lower limit, (ii) human palm, (iii) human finger, and (iv) upper limit as reported in Table 2. **(B)** Comparison of the information gained when doubling the density of the most informative class (central bar) or when combining with a different population (right bar). The baseline density (left bar) was set at 10 afferents/cm^2^.

A substantial increase in the amount of encoded information was found for all features when increasing the density from the lower limit to realistic densities. Conversely, increasing the densities from the finger values further to the upper limit did not lead to additional increases of encoded information, neither when considering individual classes nor their combination, suggesting that the information in multi-class population coding saturates similarly to single-class coding. The only exception was probe size, for which information content at the upper limit was higher than at the finger density. As shown in Figure 5B and summarized in Table 1, stimulus size is also the only feature for which increasing the density of the most informative class, SA1, improves the information content more than combining different classes. As discussed previously, stimulus size is a purely spatial feature, and a high density of afferents is necessary to discern small differences in the shape of the stimulus. In contrast, for all other features considered, combining the content of the two most informative afferent classes yields more information than doubling the afferent density of the most informative class alone.

**Table 1.** Information maximizing encoding strategies for each stimulus feature trading off increases in innervation density for a single afferent class versus adding fibers from a different class.

|  | Stimulus size | Frequency | Ramp length | Ramp indentation |
| --- | --- | --- | --- | --- |
| **Most informative class** | SA1 | PC | PC | SA1 |
| **Optimal encoding strategy** | Increase density of SA1 population | Add SA1 population | Add SA1 population | Add RA population |
| **Underlying rationale** | Highest gain in information by increasing SA1 density<br>PC provide no information | SA1 adds more information than RA<br>PC information is density independent | SA1 adds more information than RA<br>PC information is density independent | SA1+RA reaches the highest information content<br>PC provide no information |

Together, these results show the advantages for information encoding at the population level of spreading information across classes of receptors with complementary information rather than simply packing more receptors of a given class into the skin, even if receptors of this class are highly informative about the stimulus.

### Spatial and temporal contributions to information coding

After establishing how much information is encoded in each afferent class and their combination, we investigated the nature of this population coding in more detail. In particular, we asked two questions relevant to understanding the spatial and temporal organization of the population code. First, how important is the precise temporal structure of the population activity for stimulus decoding? Second, how important are differences in spatial neuron-to-neuron response profiles to decode stimulus information?

The importance of the spatial structure of the afferent population code for information coding, that is, the afferent-to-afferent difference in stimulus tuning properties at different spatial locations, is supported by the finding that natural tactile stimuli elicit specific firing patterns in afferents located in different places (16, 28). A critical role for the temporal structure of individual afferent activity has been demonstrated in previous studies (28, 29) and is also supported by the fact that thalamic and cortical somatosensory neurons also encode tactile information with millisecond-scale spike timing precision (30–33). However, it is unknown whether these expectations would hold at the level of afferent population coding. For example, precise spike timing might be less important when considering a full population of afferents rather than a single one. Furthermore, information in the spatial and the temporal structure might be redundant, such that information in the spatial structure could be recovered from the temporal structure or vice versa. Addressing these questions therefore requires a direct test with a large population. First, we evaluated whether the distribution of afferents in the space, parameterized in the current setup as the distance of the afferent location from the stimulation site, impacts the population coding capabilities. To do so, we repeated the analysis pipeline used to compute the information content in the spiking activity after corrupting the spatial information in the spike matrices by randomly permuting the identity of afferents. We called this “space-coding neglected” information. The difference between the original and the space-coding neglected information quantifies how much of the original information can only be accessed using the spatial structure of the code. Note that this quantification is performed at a fixed spatial density, and it is thus different from the previous analyses of the effect of spatial density. We found that, after destroying the spatial structure, information content dropped by on average 26% (Figure 6). The loss of information was higher for SA1s (39%) compared with RAs (18%) or PCs (16%).

**Fig. 6.**
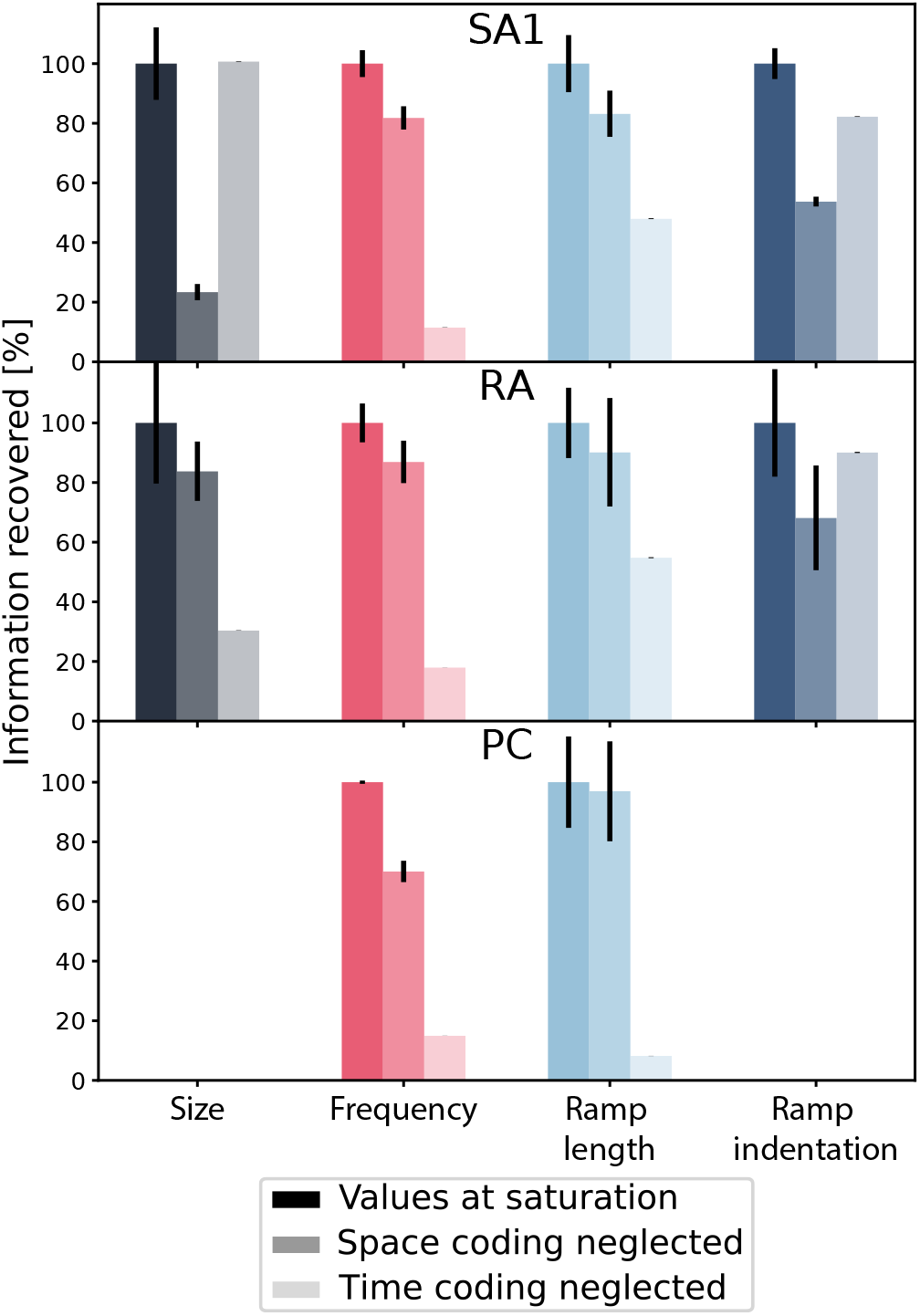
Information recovered after corrupting spatial or temporal coding. Information recovered for each afferent class after neglecting either spatial organization or precise spike timing, normalized with respect to the maximum information content in the original data at the saturation level (see Figure 3C). Note that for both stimulus size and ramp indentation, information carried by PC class was null (see Figure 3C) and such class was excluded from this analysis.

Next, to quantify the specific contribution of temporal structure to the total information in the population code, we computed a “time-coding neglected” information value from the population responses. To do so, we randomly shifted the spikes with a Gaussian distribution of zero mean and standard deviation of 10 ms, before proceeding with the rest of the analysis pipeline. The difference between the original and the time-coding neglected information quantifies how much of the original information can only be accessed using the millisecond-scale temporal structure of the code. We found that, after destroying the temporal structure, information content dropped by on average 54% across all classes (Figure 6). Information loss was highest for PCs (88%) and relatively lower for RAs (51%) and SA1s (39%). Notably, across our set of stimuli we found higher information loss when neglecting the temporal resolution of spike trains rather than the spatial distribution. This result indicates that even in large afferent populations spike timing with high temporal precision remains an important part of the neural code. In contrast, the ramp indentation for both SA1 and RA classes and the stimulus size for SA1 rely more on spatial than temporal activation, which can be explained by the spatial nature of these two features.

## Discussion

This study takes a novel perspective on stimulus coding by tactile afferent populations. Much of our current understanding of encoding mechanisms of tactile stimuli derives from electrophysiological studies. However, these are severely limited in the number of afferents that can be recorded at a time. In addition, many previous studies have focused only on those afferents terminating directly at the stimulus contact location. Thus, a biased picture of tactile coding might have emerged. In fact, to our knowledge, population coding of tactile afferents, taken as the spatiotemporal activation of multiple afferents belonging to one or *more* classes, has scarcely been investigated before. Here, we used a recently developed computational model that allows simulation of tactile neural responses at the population level with high accuracy. Although any putative population-level coding mechanisms derived from modeling would need to be experimentally verified, this approach allows investigating aspects of the neural code that are currently experimentally intractable and can therefore generate ideas for potential downstream decoding mechanisms.

### Single-class coding and receptor density

We first investigated how the density of afferents from a single class plays a role in the encoding process. We showed that the information content of both SA1 and RA populations increases asymptotically with afferent density until saturation. This effect was consistent for all features considered, although the specific saturation densities varied between features. This result highlights that tactile information is generally spread across a population of multiple afferents, even for features that are not explicitly spatial. Furthermore, the afferent class most informative about a tactile feature at low innervation densities might be different from the most informative class at high densities. Consequently, judging or predicting the information content of a population from recordings of single afferents only might be misleading and provide a biased picture of how information is represented in full populations.

In contrast to SA1 and RA afferents, the information level for PC afferents was essentially constant for all density values considered across all tactile features. While this result might be taken to suggest that the PC population does not contribute information above that of a single afferent, there is evidence to suggest that PC populations might be important in different tactile contexts than the ones explored here: making contact with surfaces causes mechanical waves to spread throughout the hand, activating PC afferents as far away as the palm and their joint population activity carries information about how contact is made and other aspects of the grasp (34).

It should be noted that for all afferent classes, the minimum density needed to recover the maximum information for any tactile feature is lower than the empirical afferent densities estimated for the human hand (11). We speculate that the minimum density of afferents required to reach the information saturation might be higher for more complex features. Indeed, as an initial investigation into the power of largescale neural simulations on a population level, this study considered relatively simple stimulus features compared to the complexity of realistic tactile interaction. Similarly, previous studies showed a strong relationship between SA1 density and tactile spatial acuity (11): afferents, particularly of SA1 type, need to be densely packed in the skin to resolve and discriminate extremely fine features. While our setup included one clearly spatial stimulus (probe size), none of the others were purely spatial. Finally, afferent innervation densities across most of the skin of the human body are much lower than those in the hand and indeed within the range identified in the current study, suggesting that our stimulus set was covering a large part of the physiologically relevant range.

Interestingly, we found that the RA class at saturation density tended to encode less information than the SA1 and PC populations, but in contrast, was more informative than SA1s at low densities (<10 afferents/cm^2^). This result suggests that the way information is spread across afferent classes depends in part on receptor density, and in turn, should affect optimal decoding downstream. Indeed, tactile innervation density changes dramatically across different body areas, both in terms of the absolute number of afferents and relative innervation densities of different classes (11), and it is possible that changes in the class composition at different skin sites partially reflect density-dependent optimal allocation of afferent classes. Our findings also suggest that tactile information need not be linked firmly to a given receptor type, but that information is spread in a dynamic way across different afferent classes (see 19, for a concrete example in frequency coding).

### Complementarity and redundancy across afferent classes

The second step of our analysis was to consider combinations of afferent classes and to evaluate their information content with respect to different stimulus features. Here, we found both redundant and complementary contributions to the information across afferent classes. All afferent classes generally provided at least some complementary information about stimulus features, suggesting that downstream areas should integrate information from different classes to maximize information (see also 15). Quantifying such complementary information is a necessary first step towards further study of submodality convergence in the stimulus encoding process, especially considering that directly accessing the integration mechanisms in humans is complicated. Convergence has previously been inferred from cortical recordings in primates for multiple individual stimulus attributes (18, 35, 36) before, but here we quantitatively demonstrate that information is spread across afferent types in most cases, and therefore, submodality integration can be expected to be a general feature of downstream processing. Not all information was complementary however, and we also found considerable degrees of redundancy between afferent classes. Redundancy in neural coding has been extensively debated (see 37, for a review) and can be a strategy for robust stimulus encoding. Indeed, over-representing stimulus information using large populations of neurons increases the probability of having a relevant impact in downstream neurons, guaranteeing —or, at least, making more plausible— that critical information is processed while negligible information is discarded —or less likely used—. Redundancy can rank information according to relevance, overcoming the associated coding inefficiency in favor of a significant performance increase (38). Furthermore, redundancy could be interpreted as a strategy to make the neural code resilient in the event of temporary or permanent lack, shortage, or failure of input from an afferent class. This theory is supported by recent findings in an experimental study in mice that showed that the use of genetic ablation strategies to suppress the response of either rapidly or slowly adapting afferents leaves responses in the somatosensory cortex mostly unchanged (17), which implies that the required information can be recovered from the remaining afferent input. This would not have been possible if the two classes had encoded complementary information only. Such a process might be beneficial when several features are processed simultaneously, and redundancy between classes might help to disambiguate the stimulus.

### Information maximizing receptor selection

We investigated whether increasing the density of afferents of a given class or combining them with afferents of a different class yields higher information gain. We found that adding afferents belonging to a different class was generally more efficient than increasing the density of the most informative class by the same amount, confirming that the information about stimulus features is not segregated in single afferent classes, but is spread across them. Indeed, while absolute tactile innervation densities vary widely across the body, the fraction of slowly adapting afferents at any given site varies only between 40 and 70% and is relatively evenly split for most body regions (11), especially for those with lower innervation. Our results suggest that such a composition increases information transmission, while minimizing fiber count. The number of tactile fibers that can fit into the nerves and spinal cord is naturally constrained, and consequently, extensive skin areas are innervated at low density. Neurons are also energetically expensive, and therefore it is plausible that evolutionary optimization might have maximized the ratio between information and energy consumption by spatially distributing the mechanoreceptors and diversifying response properties across different receptor classes.

In several sensory systems other than touch neural populations are also composed of multiple cell classes with distinct response properties. Indeed, early sensory pathways frequently split into different classes with disparate response properties (6, e.g. the large number of retinal ganglion cell classes). According to the efficient coding hypothesis, sensory systems have evolved to optimally transmit information about the surrounding world, given constraints on their biophysical components and energy use (39). This theory also explains splitting a population into two or more cell classes as a strategy to maximize information transmission, as shown in previous studies for different sensory systems (8–10). Our findings support this hypothesis, showing that, in most cases, a combination of classes was more informative than a single class higher-density population.

### Limitations and future work

Our study focused on the three main classes of tactile afferents that mediate discriminative touch, but other classes, such as SA2 afferents that primarily signal skin stretch or unmyelinated afferents, also contribute to tactile coding. Furthermore, tactile innervation and neural response properties differ somewhat in the hairy skin (11), which covers most of the human body. Thus, our results will most directly reflect tactile coding on the human hand, but future studies should consider how these results might extend to other regions of the body.

As the findings are based on computer simulations, the veracity of the results will depend on the accuracy with which the spiking responses can be replicated in the computational model. The stimuli we used, namely indentations by a single probe orthogonal to the skin surface with a superimposed vibration, are similar to those on which the original model was fit and fall into the range where it has been validated most extensively (20). Still, by combining multiple tactile features, we believe that our simulated stimuli are sufficiently complex, varied, and natural that the resulting findings can be considered of behavioral importance. One avenue for future research would be to investigate information transfer on tactile inputs arising from natural behaviors such as grasping and manipulating objects, which include multiple contacts, shear forces, and movement between the object and the skin. However, this would require further work on the precise spatiotemporal force patterns on the hand during such behaviors and spiking models that take into account more complex afferent response properties (see 40, for an example).

To study the effects of different innervation densities, we considered a simplified setup, distributing the afferents over a single dimension while neglecting some properties affecting the spatial distribution of afferents, for example the complex shape of the human hand. Future studies should take this aspect into account to reveal how the shape of the hand, the different afferent densities, and the composition of the population in different areas of the hand play a role in stimulus encoding. In the same direction, population coding strategies and afferent distribution might be coupled with natural stimulus statistics in different body areas to deepen the understanding of how the human somatosensory system is optimised to receive and process natural tactile stimuli.

## Materials and Methods

### Simulation of spiking activity

To generate the spiking activity of tactile afferents, we used a previously published and validated model called Touchsim (20). We employed the model to simulate populations of SA1, RA, and PC afferents terminating along a line of 1 cm for SA1s and RAs, and 5 cm for PCs radiating outwards from the stimulus location. The density of afferents varied between 1 and 140 afferents/cm^2^ for a total of 16 different populations per afferent type. This range includes the physiological innervation densities estimated for the human hand (25). In some analyses we also directly set individual class densities to those of the human palm or finger (see Table 2 for precise values).

**Table 2.**
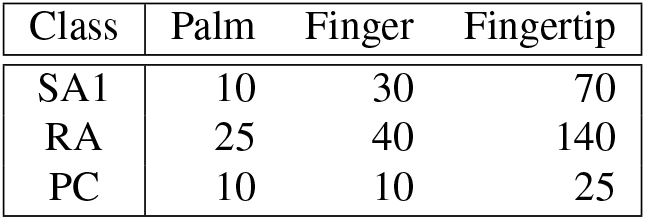
Estimated innervation densities of afferent classes (afferents/cm^2^) for different regions of the human hand (25).

| Class | Palm | Finger | Fingertip |
| --- | --- | --- | --- |
| SA1 | 10 | 30 | 70 |
| RA | 25 | 40 | 140 |
| PC | 10 | 10 | 25 |

We designed stimuli with circular shapes, which are indented in the skin following a ramp-and-hold function (see Figure 1 B). When the maximum indentation of the ramp is reached, a sinusoidal wave is superimposed. This setup simulates well-established psychophysical setups in which a probe is brought into contact with the skin and then vibrated at a set frequency. It also includes many aspects of natural tactile stimulation: indentation, retraction, and constant stimulation at different depths and spatial scales, as well as vibrations at different frequencies. Individual stimuli are created by varying 4 different features: 1) the stimulus size (4 conditions: [1:1:4] mm), 2) the maximum indentation (4 conditions: [0.3:0.3:1.2] mm), 3) the ramp-up time (5 conditions: [0.01:0.01:0.05] s), and 4) the frequency of the superimposed sinusoidal wave (10 conditions: [0, 10, 20, 40, 60, 80, 100, 130, 160, 200] Hz). This setup yielded 800 unique stimuli, and the afferent response to each was simulated for 40 trials. The model included simulated neural noise. Additionally, in order to simulate environmental noise such as motor noise during active touch, we jittered the stimulus location (by *±* 0.3 mm), the amplitude of the sinusoidal wave (by *±* 0.05 mm), and the ramp indentation (by *±* 0.1 mm) on every trial.

### Dimensionality reduction

Information calculations from high dimensional data require prohibitively large datasets. A common strategy to address this issue is by performing dimensionality reduction on the data. Here, we used Non-Negative Matrix Factorization (NMF) to decompose the spatio-temporal matrix of spiking responses across the population.

Responses were discretized by binning the spike trains into 2 ms long intervals and counting the number of spikes falling into each bin. This resulted in a matrix *R* ∈ **R**^*M* × *TN*^, where *M* is the total number of trials, *T* the number of time bins, and *N* the number of afferents in the population. NMF decompositions are naturally suited to describe spatio-temporal matrices of spiking responses, because spike trains are nonnegative, and because often recurrent spike patterns may be non-orthogonal (as NMF) and partly overlapping (explainable by the same underlying activity pattern). NMF describes a single trial spike train as a sum of trial-independent non-negative spatiotemporal modules (describing the most often recurring spatio-temporal firing patterns) and trial-dependent non-negative activation coefficients representing the strength of recruitment of each module in the considered trial (22, 23):

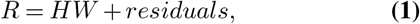

where *H* ∈ **R**^*M* × *K*^ contains the non-negative activation coefficients for the *K* modules in each trial and *W* ∈ **R**^*K* × *TN*^ contains the non-negative modules. We used the function *NMF* included in the scikit-learn Python library (41) to calculate the NMF decomposition.

We performed the NMF decomposition separately for each of the three afferent classes at each density value considered. Beforehand, we randomly separated the whole set of trials into balanced sets with a 25/75 split. We used the 25-set to determine the number of modules *K* as the minimum number of modules capable of explaining a selected level of variance of the original data in *R*, as follows. First, to consistently select the level of variance explained between populations of the same class but with different densities, we calculated the saturation level of the accounted variance for each population considered (tolerance <1%). We averaged the saturation levels across populations of the same class with different densities and used this value as the new threshold for the explained variance. Finally, we calculated *k* modules *W* and activation coefficients *H* on the same 25-set. Given the *W* modules from the 25-set, we computed the activation coefficients *H* on the remaining 75-set.

### Stimulus decoder

After dimensionality reduction, we fed the activation coefficients *H* computed with the NMF to a stimulus decoder. We used multinomial logistic regression to decode each stimulus feature separately on a trial-by-trial basis based on the neural activity (similarly to (24)). The scikit-learn Python library (41) was used for the implementation. This type of classifier uses a linear function *f* (*s, i*) to predict the probability of outcome *s* for trial *i* such that:

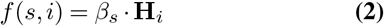

where **H**_*i*_ is a vector containing the NMF activation coefficients for trial *i* and *β*_*k*_ stores the coefficients associated with outcome *s*. When generalizing to *S*_*n*_ features, the multinomial logistic regression model consists of *S*_*n*_ − 1 independent logistic regression models regressed against the remaining *S*_*n*_ outcomes. Note that outcomes correspond to the possible values that the stimulus features could take and vary for each feature.

The 75-set was divided equally and stratified into training and test sets. We trained the classifier on the activation coefficients of the training set and evaluated performance using the activation coefficients of the test set. The training procedure was performed using a stratified 5-fold cross-validation. This process was repeated for each population of afferents (both for single and combined classes) and all afferent densities. The solver used for the fitting procedure was *lbfgs* in combination with *L2* regularization. We selected the parameter *C* for the regularization by performing grid search. The scoring of the classifier was the negative log-likelihood, also known as the cross-entropy loss.

The final fitted model outputs the posterior probability of observing each stimulus feature given the neural activity captured in the NMF activation coefficients (24). From this posterior probability, we decoded the stimulus *ŝ* that was most likely given the observed afferent activity.

### Mutual Information

Next, we computed the mutual information (42) from the confusion matrix of the decoder as follows (2):

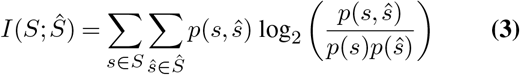

where *S, Ŝ* stand for the set of all possible presented and decoded stimuli, respectively. *p*(*s, ŝ*) denotes joint probability distribution, which is derived from the confusion matrix obtained empirically across all trials, of presenting stimulus *s* and decoding stimulus *ŝ* in a given trial. *p*(*s*) and *p*(*ŝ*) correspond to the marginal probabilities of *s* and *ŝ*, respectively. The information in the confusion matrix is a data-robust lower bound to the total information carried by population activity. This approximation is tight when neural activity can be categorically binned into as many values as the number of distinct stimuli without losing considerable information. The information in the confusion matrix captures aspects of information processing, such as the distribution of decoding errors, which are not captured by simple measures such as the fraction of correctly decoded stimuli (2). Since the information upper bound is the entropy of the stimulus set (indicating perfect single-trial stimulus discrimination), we normalised information values by dividing them by the entropy of the stimulus set:

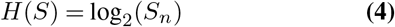

where *S*_*n*_ is the number of values that the stimulus can take.

### Computation of complementary information

To assess the complementarity of stimulus information carried by different classes, we computed the information carried about each stimulus feature by the joint activity of populations of two or three afferent classes and compared it to the information carried by a single-class population. We defined the amount of information carried by the pair of afferent classes that is complementary to that of a reference class as the difference between the information carried by all the classes (including the reference class and the additional ones) and the information carried by the reference class. We repeated this process, taking each class as the reference class in turn. As an example, for SA1 afferents as the reference class, the complementary information is computed as:

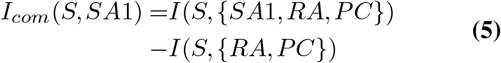

We defined the redundant information between the additional classes and the reference class as the sum of the information carried individually by the reference class and the additional ones minus the information carried by all the classes together, such that (again, taking SA1s as the reference class):

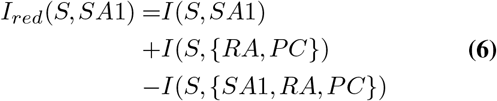

The sum of redundant (eq. 6) and complementary (eq. 5) information for a class equals the total information carried by that class.

## ACKNOWLEDGEMENTS

This work was supported by the EU Horizon 2020 research and innovation programme under grant agreement 813713 (NeuTouch).

---

**Fig. S1.**
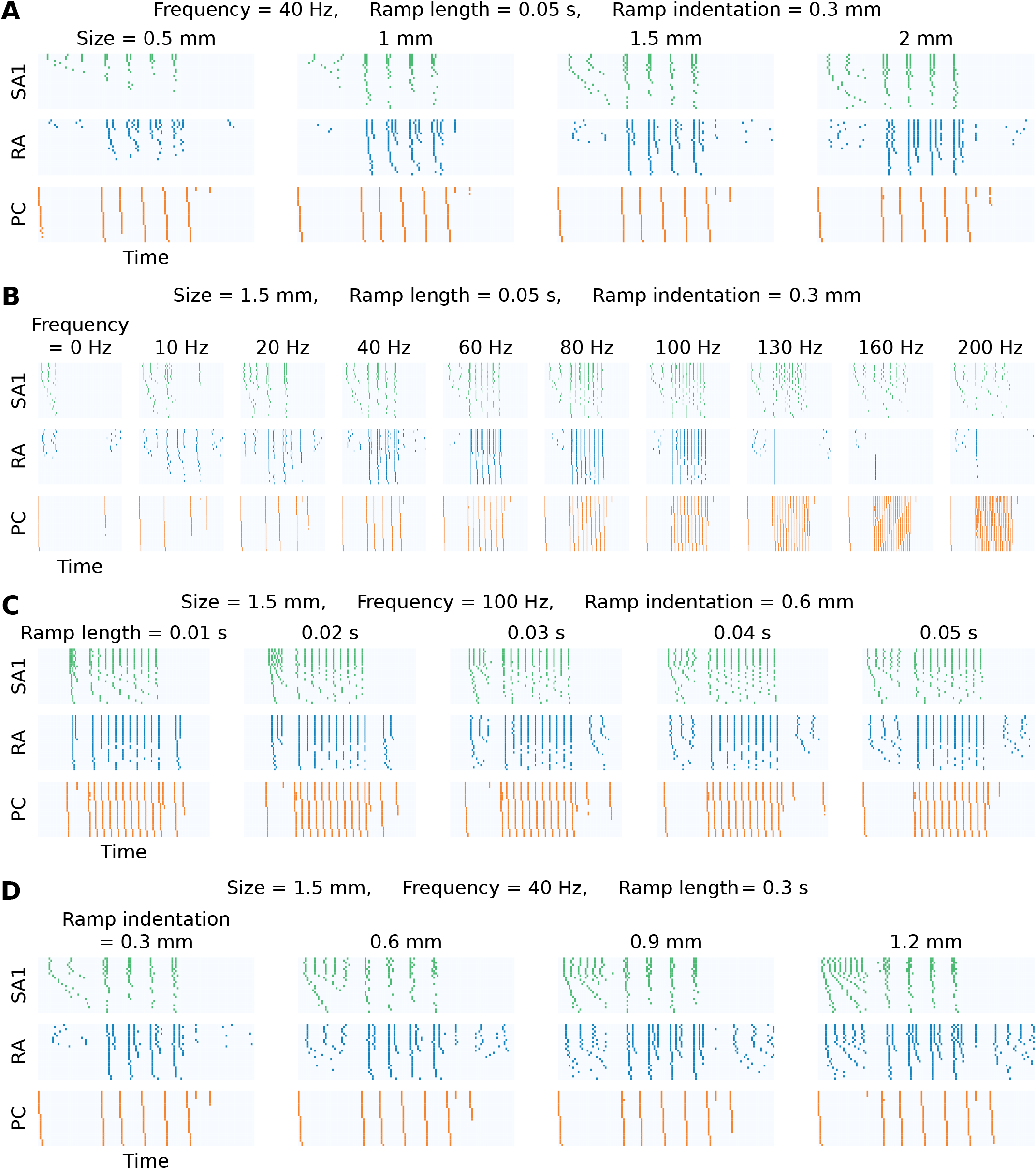
Illustrative examples of simulated spiking activity for the three afferent classes as a function of **(A)** stimulus size, **(B)** frequency, **(C)** ramp length, and **(D)** ramp indentation depth. Note that we have conditioned on the remaining features for each panel and that the afferent densities chosen in this example correspond to the ones in the finger. Note that the spike trains are shown by conditioning on the remaining variables.

